# Novel plasmalogen derivative KIT-13 restores neurological function in a mouse model of Rett syndrome by reducing neuroinflammation and restoring mitochondrial function

**DOI:** 10.1101/2025.06.11.656246

**Authors:** Akane Matsuda, Hideyuki Nakashima, Takehiko Fujino, Tatsuo Okauchi, Md Shamim Hossain, Yasunari Sakai, Shouichi Ohga, Kinichi Nakashima, Masanori Honsho

**Author notes:** These authors contributed equally to this work (Akane Matsuda, Hideyuki Nakashima). These authors are co-corresponding authors.

## Abstract

Neurodevelopmental disorders, including Rett Syndrome (RTT), have no functional cure and cause substantial levels of disability. Neuroinflammation is now strongly associated with both neurodevelopmental disorders and neurodegenerative diseases, providing a strong rationale for development of novel therapeutics targeting common neuroinflammatory mechanisms. RTT is caused by mutations in the methylated DNA binding factor *MECP2. Mecp2*-deficient (*Mecp2*-KO) mice, which have been extensively characterized as a mouse model of RTT, exhibit high levels of neuroinflammation, mitochondrial dysfunction, and severe neurological symptoms similar to RTT patients. KIT-13 is a novel plasmalogen derivative being developed for the treatment of neurodevelopmental disorders including RTT. This study evaluated KIT-13 in both cell-based and *in vivo* models for its potential to inhibit neuroinflammation and address underlying mitochondrial dysfunction, as well as effects on RTT-like neurological symptoms in the RTT mouse model. Oral administration of KIT-13 to *Mecp2*-KO mice significantly reduced neurological symptoms assessed by a composite score evaluating mobility, gait, hindlimb clasping, tremor, breathing, and general condition and improved the life span of the RTT model mice. In addition, KIT-13 suppressed mitochondrial DNA leakage associated with *Mecp2* deficiency, and significantly suppressed neuroinflammation as measured by microglial cell morphology. These results suggest that KIT-13 may be a promising therapeutic agent for RTT and other neuroinflammation-related diseases.

## Introduction

Neurodevelopmental disorders (NDDs) include a wide range of neurological conditions that begin in childhood and disrupt the proper development and function of the brain and nervous system; NDDs are prevalent, affecting approximately 15% of children and adolescents worldwide (1, 2). Most NDDs are the result of genetic mutations that manifest early in development, with tremendous impact on physiological, intellectual, emotional, and social development. Gene therapy approaches are still in very early stages of clinical development, and symptomatic treatments available today are limited to anxiety, impulsivity, and some other symptoms (3,4). There is a significant need for new disease modifying and improved symptomatic treatments, warranting the development of more effective treatments.

Previously, NDDs were thought to result solely from abnormal function of neuronal cells, however recent studies have demonstrated that glial cells may also be involved in the underlying pathophysiology of NDDs (5,6). In fact, microglial hyperactivation has now been reported in neurodegenerative diseases such as Alzheimer’s disease (7) and in NDDs including autism spectrum disorder (ASD) and Rett syndrome (RTT) (8,9). Both neuroinflammation and mitochondrial dysfunction are observed concomitantly in approximately 65% of RTT patients (9). Mitochondrial dysfunction results in loss of membrane integrity and leakage of mitochondrial DNA (mtDNA) into the cytoplasm (10), which triggers neuroinflammation through the activation of DNA sensing receptors (11). Therefore, it is reasonable to hypothesize that mitochondrial dysfunction is also one of the major pathogenetic causes of neuroinflammation in NDDs, including RTT.

Plasmalogens are a subclass of glycerophospholipids which are abundant in the brain and have neuroinflammation inhibitory properties (12,13). Plasmalogens have been shown to be decreased in patients suffering from neurodegenerative diseases including Alzheimer’s disease (14) and in several neurodevelopmental disorders including RTT (15,16), leading to the hypothesis that increasing plasmalogen levels in these disorders could provide important therapeutic benefit. Oral administration of natural plasmalogen extracts in clinical studies has been reported to alleviate clinical symptoms of mild Alzheimer’s disease and Parkinson’s disease (17,18), however efficacy of oral treatments may be limited by the low stability of natural plasmalogens under acidic conditions, such as in the gastric compartment, and large-scale production of plasmalogen extracts may be limited by availability of natural source materials. The development of novel plasmalogen derivatives with greater biological and chemical stability and more drug-like pharmacokinetic properties which would enhance tissue biodistribution may provide compounds with increased therapeutic efficacy and stronger anti-neuroinflammatory effects than natural plasmalogens (19).

RTT is a neurodevelopmental disorder caused by mutations of the methylated DNA binding protein *MECP2. Mecp2*-knockout (KO) mice have been developed and are well characterized as a model of RTT in which many aspects of RTT symptoms are recapitulated and neuroinflammation pathology is present. In this study, a panel of newly developed plasmalogen derivatives called ‘KIT’ compounds were evaluated in *Mecp2*-KO mice for potential efficacy in reducing RTT symptoms as a representative NDD model characterized by neuroinflammation-based pathophysiology. Among the plasmalogen derivatives tested, KIT-13 demonstrated the most robust improvements in increasing survival and reducing RTT-associated neurological symptoms in *Mecp2*-KO mice. In addition, treatment with KIT-13 showed anti-neuroinflammatory activity in the *Mecp2-*KO mice, and potently inhibited DNA leakage from mitochondria which is believed to be associated with *Mecp2* deficiency and an underlying cause of neuroinflammation in RTT. Together, these results support further development of KIT-13 as a potential drug treatment for RTT with the potential for therapeutic efficacy in additional neurodevelopmental disorders.

## Results

### KIT-13 improves neurological symptoms and restores healthy microglia morphology in RTT mice brain tissue

With the aim to develop highly effective therapeutic agents for the treatment of NDDs with underlying neuroinflammation, novel glycerophospholipids were designed by modifying the chemical structure of natural plasmalogen, which is known to have anti-neuroinflammatory activity. Four selected plasmalogen derivatives were synthesized (KIT-8, KIT-13, KIT-19, and KIT-20, shown in Extended data Fig. 1a), which have optimized drug-like properties compared to natural plasmalogen. The therapeutic effects of these four plasmalogen derivatives were tested, in parallel to a natural plasmalogen extract, in *Mecp2*-KO mice, a well-characterized mouse model of RTT created by a genetic deletion in the *Mecp2* gene (20,21). Key outcome measures were survival rate, a composite neurological symptom score for RTT-like symptoms (22) and body weight. With oral administration over a period of six weeks, KIT-13 was the most effective of the treatments tested. KIT-13 significantly improved the survival of *Mecp2*-KO mice (Fig. 1a) and reduced the RTT neurological symptom score including improvement of the general condition of the *Mecp2*-KO mice (Fig. 1b). KIT-13 effects were greater than effects seen with the natural plasmalogen extract, and both RTT symptom scores and survival increases associated with KIT-13 were consistently better than the three other plasmalogen derivatives tested. No effects on body weight were observed in *Mecp2*-KO mice following oral administration of KIT-13 (Extended data Fig. 1b). These results suggested that KIT-13 may be particularly well suited for development as a therapeutic agent for RTT, and further investigation was warranted to characterize its mechanism of action in this mouse model.

**Fig. 1.**
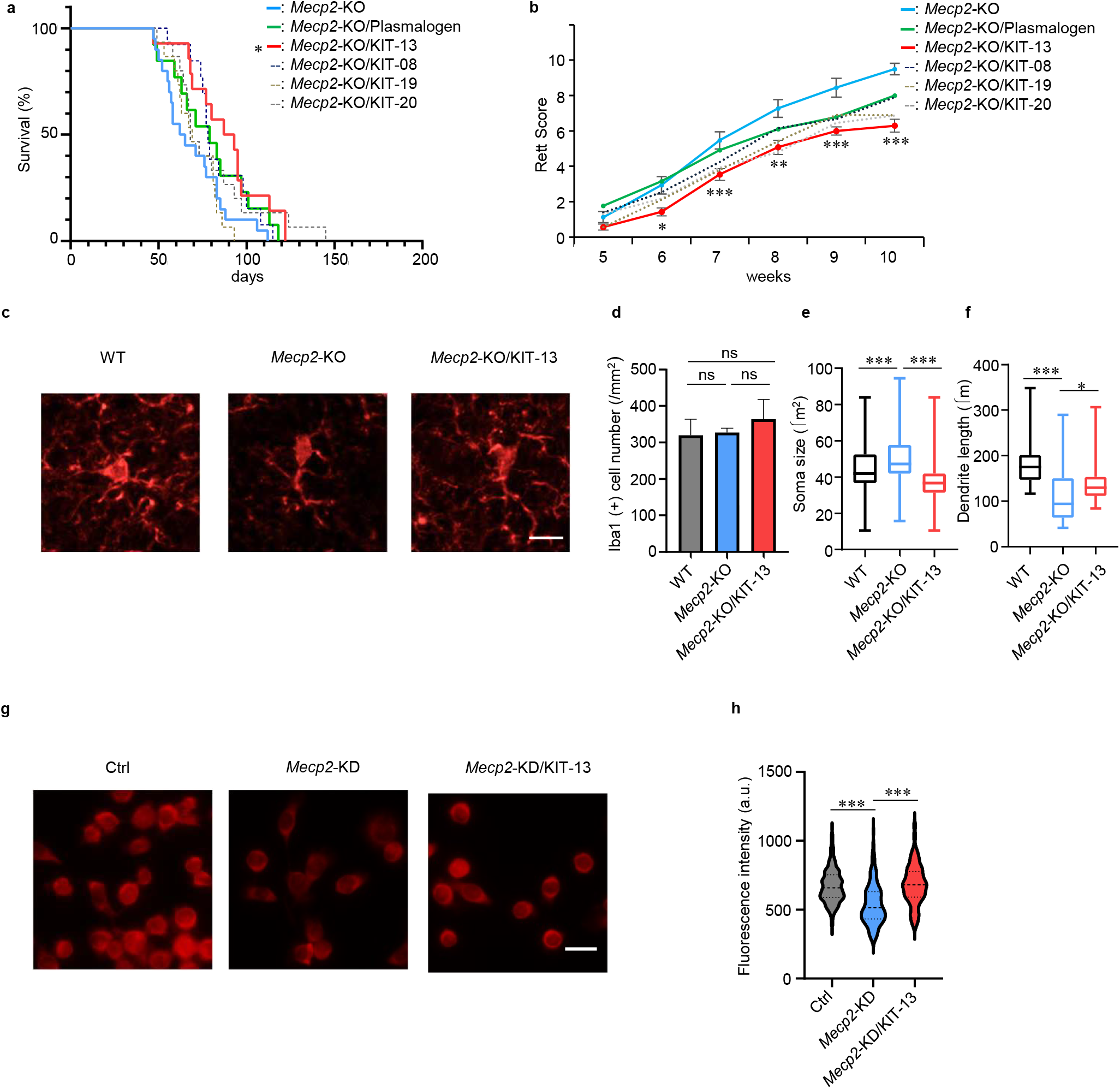
KIT-13 is effective for the general condition of RTT model mice and dysregulated microglia. **a**, Kaplan-Meier curves of *Mecp2*-KO (n = 20), KIT-13-treated *Mecp2*-KO (n = 14), and *Mecp2*-KO mice treated with other novel plasmalogen derivatives. ^*^, *P* <0.05; versus *Mecp2*-KO mice (Log-rank test). **b**, Time plot scores from neurological symptoms assessment of *Mecp2*-KO (n = 15), KIT-13-treated *Mecp2*-KO mice (n = 14) and other plasmalogen derivative-treated *Mecp2*-KO mice. Data are represented as mean ± SEM. The time plot of neurological symptom scores is shown after one week of KIT or plasmalogen ingestion at a dose of 20 μg/kg/day through drinking water, starting at 4 weeks of age. Wilcoxon rank-sum test with normal approximation was performed at each time point to compare the *Mecp2*-KO group and the KIT-13-treated *Mecp2*-KO group. ^*^*P* <0.05, ^**^*P* <0.01, ^***^*P* <0.001. **c**, Representative immunofluorescence images of wild-type (WT), *Mecp2*-KO, and KIT-13-treated *Mecp2*-KO mice microglia in the hippocampus. Scale bar = 10 μm. **d-f**, Quantitative assessment of Iba1-positive microglial cell number (**d**), soma size (**e**), and total dendrite length (**f**). Microglia were measured from n = 3 mice per group. Statistical testing was performed using Tukey’s multiple comparisons test. ^*^*P* <0.05, ^***^*P* <0.001. **g**, Representative immunofluorescence images of control (Ctrl), *Mecp2*-KD, and KIT-13-treated *Mecp2*-KD BV2 cells. Scale bar = 20 μm. **h**, Restoration of mitochondrial membrane potential by KIT-13. The reduced mitochondrial membrane potential of *Mecp2*-KD BV2 cells (center) was restored by KIT-13 treatment (right). Tukey’s multiple comparisons test. ^***^*P* <0.001.

A possible mechanism of action underlying the effects of KIT-13 in *Mecp2*-KO mice is a decrease in the level of overactivated microglia in the brain. Microglia are brain-resident immune cells which are strongly implicated in neuroinflammation processes (23). Abnormalities in microglia including larger cell bodies and shorter projection lengths have been reported in multiple brain regions of *Mecp2*-KO mice (24). Abnormalities in brain microglial cells were observed in the vehicle treated *Mecp2*-KO mice compared to WT, consistent with previous reports, and treatment with KIT-13 was associated with normalized glial cell morphology (Fig. 1c). Quantitation of effects, shown in Figure 1d-f, demonstrates that oral administration of KIT-13 significantly reduced the soma size and increased the dendrite length in the *Mecp2*-KO mice, partially restoring a healthy phenotype. Furthermore, similar results were obtained in the frontal lobe (Extended data Fig. 1c-f). Taken together, these data demonstrated that KIT-13 has an anti-neuroinflammatory effect in the *Mecp2*-KO mouse brain which may constitute part of the underlying mechanism of action of KIT-13 in reducing RTT-like neurological symptoms and in increasing survival. These results also suggest that KIT-13 is likely the best candidate among the four plasmalogen derivatives tested for further development as a potential therapy for RTT.

### KIT-13 ameliorates mitochondrial functional deficits induced by *Mecp2* gene deletion

As mentioned above, it has been reported that mitochondrial dysfunction is observed in RTT patients (9), and that loss of mitochondrial membrane integrity is expected to release mtDNA that could trigger neuroinflammatory processes. To characterize mitochondrial function in the RTT mouse model, mitochondrial membrane potential was evaluated *in vitro* in BV2 cells (control cells) and in a *Mecp2*-knockdown (*Mecp2-*KD) mouse BV2 microglial cell line. *Mecp2*-KD cells showed significantly decreased mitochondrial membrane potential, and mitochondrial membrane potential was fully recovered following the treatment of these cells with KIT-13 (Fig. 1g,h). To confirm elevated levels of mtDNA in the cytosol of the *Mecp2*-KD cells, which would provide additional evidence for loss of mitochondrial membrane integrity, cytosol samples were prepared by selective permeabilization of cell membranes with digitonin and subjected to qRT-PCR analysis using primers specific for four different mtDNA-specific genes (*Mt-Rnr2, MT-Co1, Mt-Nd2, Mt-Cytb*). The relative abundance of all four mtDNA-specific genes was increased in *Mecp2*-KD cells compared to control BV2 cells, and treatment with KIT-13 significantly decreased levels of all four genes in the cytosol, consistent with KIT-13 effects in restoring mitochondrial membrane integrity (Fig. 2).

**Fig. 2.**
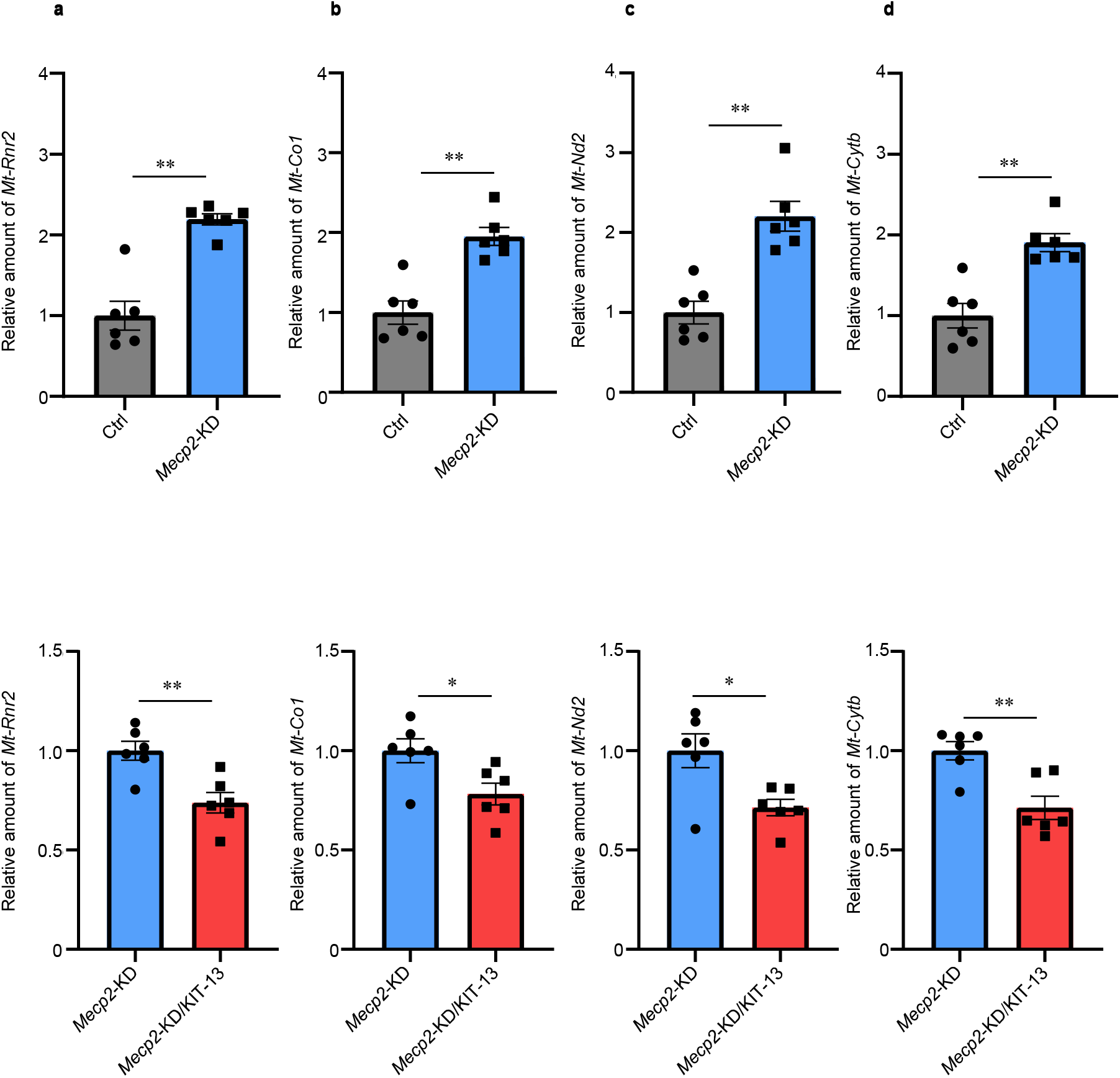
KIT-13 suppresses mitochondrial DNA leakage in *Mecp2*-KD cells. **a-d**, Recovery of mitochondrial DNA leakage by KIT-13. The increased mitochondrial DNA leakage of *Mecp2*-KD BV2 cells (top) was suppressed by KIT-13 treatment (bottom). The relative amount in the cytoplasm of *mitochondrially encoded 16S rRNA* (*Mt-Rnr2*) (a), *Cytochrome c oxidase 1* (*Mt-Co1*) (b), *NADH dehydrogenase 2* (*Mt-Nd2*) (c), and *Cytochrome b* (*Mt-Cytb*) (d). Mann-Whitney test. ^*^*P* <0.05, ^**^*P* <0.01.

Another method to evaluate mitochondrial membrane integrity is to monitor levels of the translocator protein (TSPO), an outer mitochondrial membrane protein. Increased levels of TSPO are associated with neuroinflammatory and neurodegenerative disorders (25, 26), and accordingly, TSPO has recently been used as a marker for neuroinflammation in clinical practice (27). In order to further investigate the molecular mechanism of action of KIT-13, expression levels of TSPO in *Mecp2*-KO mice were characterized with and without treatment of KIT-13. Expression of TPSO was upregulated in the cortex of untreated *Mecp2*-KO mice, and oral administration of KIT-13 was associated with decreased TSPO expression (Fig. 3a). Similarly, TSPO expression was increased in untreated *Mecp2*-KD BV2 cells and reduced after cells were treated with KIT-13 (Fig. 3b). Moreover, decreased mitochondrial membrane potential in *Mecp2*-KD BV2 cells was restored by treatment with PK11195, a TSPO receptor inhibitor (Fig. 3c). Taken together, these results suggest that KIT-13 ameliorates the impaired mitochondrial membrane potential by suppressing TSPO expression, thereby preventing mtDNA leakage from mitochondria.

**Fig. 3.**
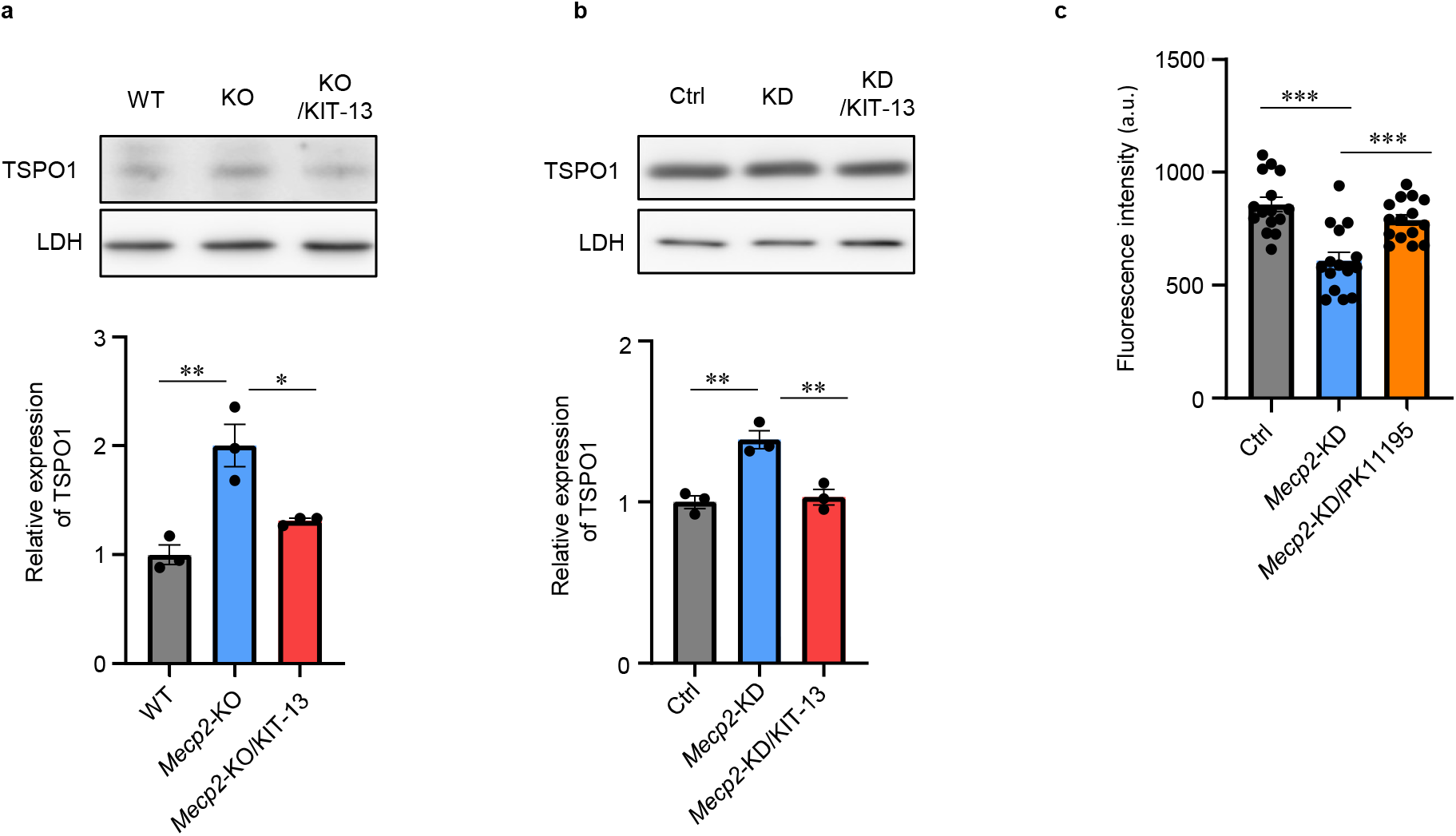
KIT-13 ameliorates the overexpression of mitochondrial membrane component protein in *Mecp2*-KD cells. **a**, KIT-13 downregulated TSPO1 protein expression in the cerebral cortex of *Mecp2*-KO mice. TSPO1 levels relative to control cells are shown (lower graph, n = 3). Dunnett’s post-hoc test. ^*^*P* <0.05, ^**^*P* <0.01. **b**, KIT-13 restored TSPO1 protein levels in *Mecp2*-KD cells to control cell levels. The TSPO1 protein expression levels of the samples were quantified and represented by the values of the amount of the TSPO1 in BV2 cells transformed with the control vector as 1. Values are averages of data from three independent experiments (mean ± SEM). Statistical significance was determined by Dunnett’s post-hoc test. ^**^*P* <0.01. **c**, The TSPO1 antagonist PK11195 reversed the reduced mitochondrial membrane potential in *Mecp2*-KD cells to control cell levels. Dunnett’s post-hoc test. ^***^*P* <0.001.

Based on these results, it is reasonable to hypothesize that mitochondrial dysfunction is one of the major pathogenetic causes of NDDs including RTT, and that KIT-13 may help restore normal mitochondrial function as a component of the mechanism of action underlying potential therapeutic benefits.

### KIT-13 attenuates abnormalities in social and anxiety-like behavior in RTT model mice

As described above, neurodevelopmental disorders such as Rett Syndrome manifest early in development with tremendous impact on quality of life. This includes the onset of physiological deficits, such as those measured by the Rett score in RTT model mice, as well as impairment of intellectual, emotional, and social development. Relatedly, RTT model mice have also been reported to show abnormalities in sociability and anxiety-like behavior, which can be measured prior to onset of the severe neurological symptoms.

To investigate the effect of KIT-13 on these aspects of behavior, *Mecp2*-KO mice at 6 weeks of age received KIT-13 solubilized in drinking water for 2 weeks, and a battery of behavioral analyses was conducted. In the elevated plus maze test, *Mecp2*-KO mice spent more time in the open arm of the maze than wild-type (WT) mice, and this was partially decreased following treatment with KIT-13 (Fig. 4a). In the three-chamber test, *Mecp2*-KO mice showed normal sociability, and this was not affected by KIT-13 treatment (Fig. 4b). Interestingly, in the three chamber-test, *Mecp2*-KO mice showed impairment in social novelty seeking, and KIT-13 restored this behavior (Fig. 4c). In the fear conditioning test, *Mecp2*-KO mice showed a decrease in the percent of time spent in freezing behavior. Treatment effects of KIT-13 did not reach statistical significance, however, *Mecp2*-KO mice treated with KIT-13 showed a trend toward normalization in the cued, but not in the contextual, fear memory test (Fig. 4d,e). Importantly, the mice in these studies exhibited minimal neurological symptoms and similar mobility at the time of the study. Mobility was measured using quantification of total distance traveled in an open field for both wild-type (WT) and *Mecp2*-KO mice (data not shown), confirming that these behavioral results were not confounded by mobility deficits. Although further investigations are required, these behavioral analyses suggest that KIT-13 may have a beneficial effect in reducing social impairment in RTT patients with *MECP2* mutations, and perhaps in other indications that are characterized by social impairment such as ASD.

**Fig. 4.**
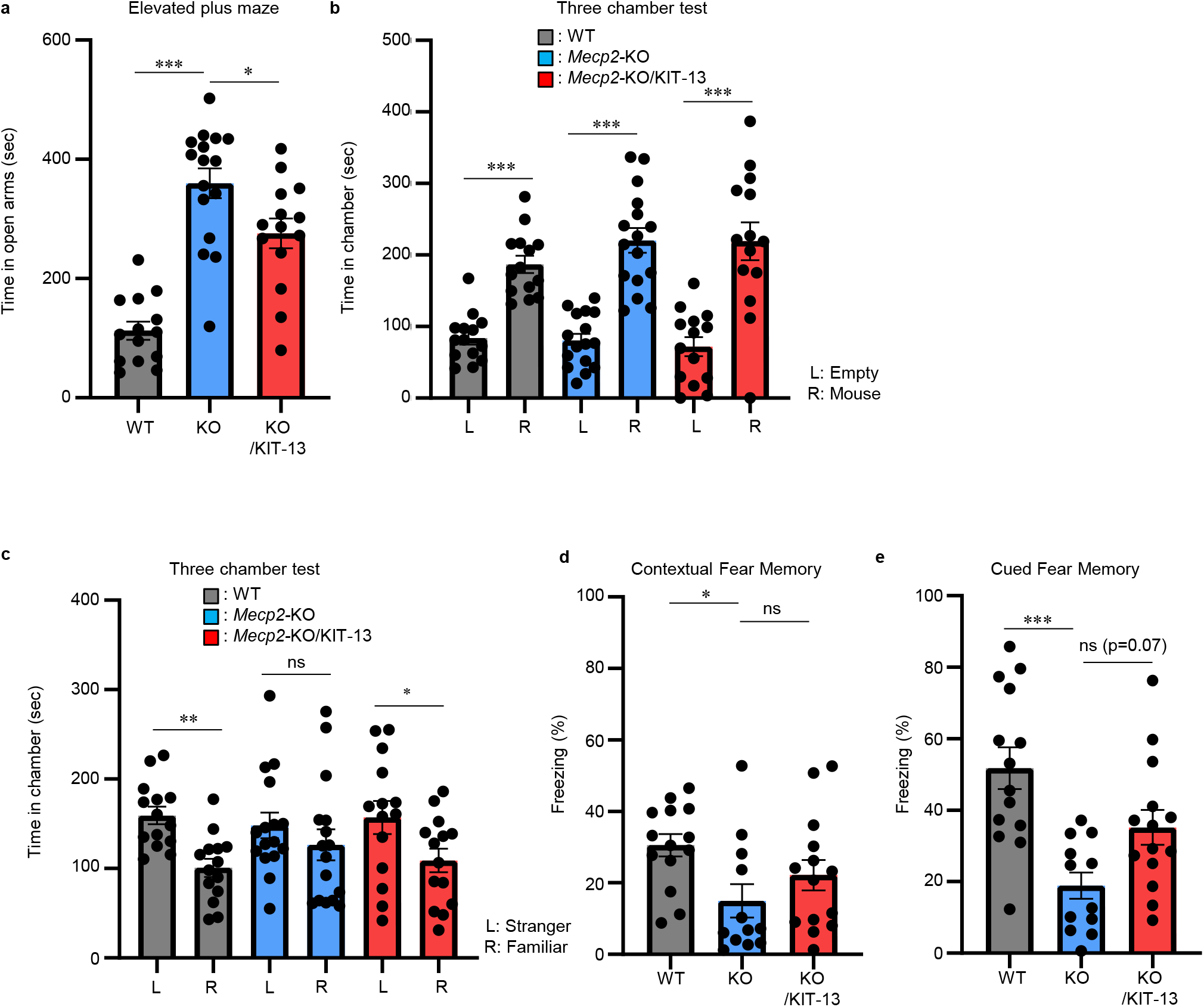
KIT-13 is effective for reducing abnormality of social behavior and anxiety-like behavior in RTT model mice. **a**, Time spent in open arms in the elevated plus maze of wild-type (WT), *Mecp2*-KO (KO), and KIT-13-treated *Mecp2*-KO mice (KO/KIT-13). Tukey’s multiple comparisons test. **b**, Sociality behavior was tested in the three-chamber assay. Student’s t-test. **c**, Social novelty was tested in the three-chamber assay. Student’s t-test. **d-e**, Quantification of freezing time in the contextual fear conditioning test (d) and the cued fear conditioning test (e). Wild-type: n = 14; *Mecp2*-KO: n = 12; KIT-13-treated *Mecp2*-KO: n = 14. Tukey’s multiple comparisons test. Wild-type: n = 14; *Mecp2*-KO: n = 16; KIT-13-treated *Mecp2*-KO: n = 14 except for the fear conditioning test. ^*^*P* <0.05; ^**^*P* <0.01; ^***^*P* <0.001; ns, not significant.

## Discussion

KIT-13 is a novel plasmalogen-derivative designed as a potential therapeutic agent for the treatment of neurodevelopmental disorders. Oral administration of KIT-13 has shown encouraging results in an *in vivo Mecp2*-KO model and an *in vitro Mecp2*-KD model of RTT described in these studies, specifically i) improving survival and reducing RTT-like neurological symptoms in *Mecp2*-KO mice, ii) ameliorating neuroinflammatory morphology of microglial cells in brain in *Mecp2*-KO mice, and iii) restoring mitochondrial function and reducing mitochondrial DNA leakage, which is hypothesized to be a key driver of neuroinflammatory processes in RTT. In addition, behavioral analyses suggest that KIT-13 may have a beneficial effect in reducing social impairment in RTT model mice.

KIT-13 is one of several plasmalogen derivatives designed and synthesized as potential therapeutic agents for neurodevelopmental disorders characterized by neuroinflammation. These studies demonstrate that KIT-13 was more effective than natural plasmalogen extract and also more effective than the closely related molecules KIT-8, KIT-19, and KIT-20 in the RTT models used here. The structure of KIT-13 is designed to have greater chemical and biological stability than natural plasmalogens, which may contribute to the observed improvements in relative efficacy. Among the KIT molecules tested, KIT-13 is unique in that it can be converted to a natural plasmalogen with a single enzymatic step. The ability of KIT-13 to act as a plasmalogen precursor may contribute directly to its robust efficacy in reducing neuroinflammation and restoring mitochondrial function, or the postulated increase in plasmalogen levels associated with KIT-13 treatment may be an adjunct mechanism of action that would further increase its efficacy in conditions such as RTT that are characterized by plasmalogen deficiency (16).

This study demonstrates that KIT-13 suppresses microglial overactivation, which is increasingly recognized as a critical target for therapeutic intervention in other NDDs in addition to RTT. Recent studies have suggested that astrocytes are also implicated in neuroinflammation in RTT (28) and moreover, it has been reported that astrocytes influence microglial activity through astrocyte-secreted factors (29,30). Considering these facts, further studies with KIT-13 are warranted to elucidate the precise mechanism of action on astrocyte-associated pathology.

New treatments are needed with improved tolerability profiles for RTT and other neurodevelopment disorders. Insulin-like growth factor-1 (IGF-1) is known to suppress microglial activation (31,32) and this is believed to be part of the mechanism of action for trofinetide. Trofinetide is composed of the tripeptide sequence Gly-Pro-Glu obtained by enzymatic cleavage of IGF-1 in the brain (33). It has been approved by the United States Food and Drug Administration (FDA) (34), but treatment is associated with diarrhea as a frequent side effect (35). Minocycline is a well-known drug that suppresses microglial activation (36), yet it has negative effects such as inhibiting bone growth in children (37) and has not been clinically applied to pediatric patients with neurodevelopmental disorders.

Based on the fact that KIT-13 can be metabolized to natural plasmalogens which are present in healthy individuals under physiological conditions, it is anticipated that KIT-13 will be well tolerated in a clinical setting. In this context, it is noteworthy that the dose of KIT-13 administered to mice in this study was equivalent, on a per-body-weight basis, to the dose of natural plasmalogens that has previously been shown to be well tolerated in clinical studies in patients with mild Alzheimer’s disease and mild cognitive impairment (17). No weight loss or other adverse effects were observed in the KIT-13-treated mice in this study. The clinical profile of KIT-13 has not yet been established, and patient studies are required to determine the efficacious dose in RTT.

In conclusion, the studies reported here demonstrate that KIT-13 has the potential for efficacy in addressing the pathophysiology of RTT, and provide evidence that further development is warranted. In support of this, KIT-13 was granted Rare Pediatric Disease Designation and Orphan Drug Designation from the FDA in March 2023 based on beneficial effects in improving survival and reducing neurological symptoms in the *Mecp2*-KO RTT model mice described here. Moreover, based on the elucidation of the KIT-13 mechanism of action reported here, KIT-13 may have potential for therapeutic benefit in other neuroinflammation-related disorders where mitochondrial DNA leakage has recently been reported, including ASD (38), Alzheimer’s disease (39), and Parkinson’s disease (40), in addition to RTT.

## Methods

### Mice

*Mecp2*-KO mice (C57BL/6 background) were obtained from Mutant Mouse Resource and Research Centers (MMRRC). All mice were maintained on a 12-hour light/dark cycle with free access to food and water. All aspects of animal care and treatment were carried out according to the guidelines of the Experimental Animal Care Committee of Kyushu University. Male *Mecp2*^tm1.1Jae^ mice were used in these studies.

### Rett score

Neurological symptoms of mice were scored as previously described (22). In brief, mice were scored blind to genotype and treatment status, focusing on mobility, gait, hindlimb clasping, tremor, breathing, and general condition. Each of the 6 symptoms was scored from 0 to 2; 0 corresponds to the symptom being absent or the same as in the WT, 1 to the symptom being present, and 2 to the symptom being severe.

### Cell culture and DNA transfection

BV2 cells were cultured in a high-glucose DMEM (Sigma) supplemented with 10% FBS (Sigma) and penicillin/streptomycin at 37 °C under 5% CO_2_. *Mecp2*-KD cells were established by transfection of a shRNA vector against mouse *Mecp2* (TRCN000304464, Sigma), followed by selection with puromycin (0.1 μg/mL). BV2 cells expressing sh*Control* (Ctrl) were likewise established by transfection of the sh*Control* vector that was generated by removing the target nucleotide sequence against mouse *Mecp2* from TRCM000304464 by PCR.

Cells were cultured in the presence of 5 μg/mL of KIT-13 together with either 1 μM of PK11195 (Abcam) for 48 h by changing the medium every 24 h.

### Antibodies

Rabbit antibody to Iba1 (019-19741) was purchased from FUJIFILM Wako. The secondary antibody, CF555 donkey anti-rabbit IgG (H+L), was purchased from Biotium.

### Immunohistochemistry

Sections were washed with PBS, permeabilized and blocked with blocking buffer at room temperature, and incubated overnight at 4°C with the primary antibody. After being washed with PBS, they were incubated for 2 hours at room temperature with a secondary antibody. Then, they were mounted on glass slides. Fluorescence images of brain sections were acquired using a Keyence fluorescence microscope BZ-X800.

### Immunofluorescence microscopy

For the assessment of mitochondrial membrane potential, cells were cultured in the presence of 50 nM MitoTracker™ Orange CMTMRos (Thermo Fisher, M7510) for 10 min. Cells were fixed with 4% paraformaldehyde in PBS for 15 min at RT, permeabilized with 1% Triton X-100 in PBS for 2 min at room temperature (RT), and incubated with DAPI for 20 min. Fluorescence images were randomly captured using a Keyence BZ-X810 fluorescent microscope (Keyence). The total fluorescence and area of each cell were measured in Hybrid Cell Count software (Keyence) and expressed as fluorescence intensity per unit area of each cell.

### Immunoblotting

Protein samples were separated by SDS-PAGE and electrotransferred to PVDF membrane (Bio-Rad). After blocking for 1 h in TBST (10 mM Tris pH 7.4, 200 mM NaCl, and 0.05% Tween-20) containing 5% non-fat dry milk, blots were subjected to immunoblotting with primary antibodies overnight at 4°C, followed by incubating with a secondary antibody for 2 h at room temperature. Immunoblots were developed with Clarity™ Western ECL Substrate (Bio-Rad) and scanned with an ImageQuant LAS 4000 mini imager (Fujifilm). The intensity of bands was quantified by Image J software (National Institutes of Health).

### qPCR of cytoplasmic mitochondrial DNA

Ctrl and *Mecp2*-KD cells were seeded at a density of 2 × 10^5^ cells per well in a 12-well dish and cultured for 48 h by changing the medium every 24 h. *Mecp2*-KD cells were likewise cultured in the presence or absence of 10 μg/mL KIT-13 for 48 h. Cells were washed three times with ice-cold PBS and incubated with 800 μL of ice-cold SHE buffer (0.25 M sucrose, 10 mM Hepes-KOH, pH 7.4, 1 mM EDTA) containing 25 μg/mL digitonin (Sigma) for 10 min on ice, harvested, and centrifuged at 2000 g for 10 min. Genomes were extracted from 100 μL of the 500 μL of the supernatant using the DNeasy Blood & Tissue Kit (Qiagen) according to the manufacturer’s instructions (41,42). Quantitative real-time RT-PCR was performed in a CFX Connect™ (Rio-Rad) using SYBR Premix Ex TaqTM II (Ti RNaseH Plus) (Takara Bio). The primers used are listed in Table 1.

**Table 1.**
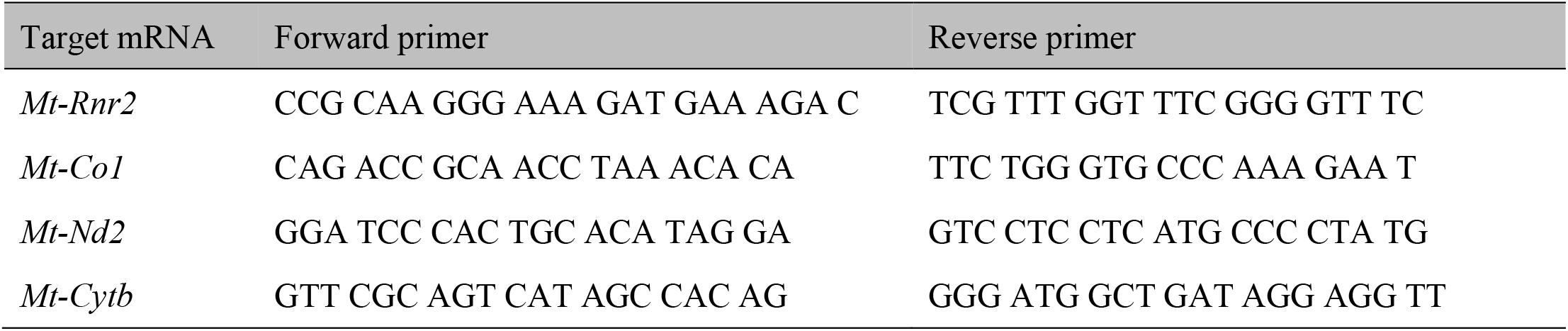
List of PCR primers used for the experiments.

### Elevated plus maze

The elevated plus maze test was performed with 6-week-old WT, *Mecp2*-KO, and KIT-13-treated *Mecp2*-KO male mice. The elevated plus maze consisted of two opposite open arms (25 cm × 5 cm × 0.25 cm) and two enclosed arms (25 cm × 5 cm × 15 cm), illuminated at 30 lux and placed 50 cm above the floor. Each mouse was placed in the central square of the maze (5 × 5 cm), facing one of the closed arms. Mouse behavior was recorded for 10 min, and the time spent on the open arm was analyzed using the automated video tracking software TimeEP1 (O’Hara & Co., Tokyo, Japan). The percentages of time spent in open arm over total time were calculated.

### Three-chamber social interaction

The three-chamber apparatus is a rectangular, non-transparent gray Plexiglas box (60 × 40 × 22 cm), partitioned into three equal-sized (20 × 40 × 22 cm) chambers by transparent Plexiglas plates with small openings (5 × 3 cm) allowing access into each chamber. One quarter-cylinder cage made of wire was placed in the corner of each of the side chambers. The box was illuminated at 20 lux. The subject mouse was placed into the central chamber and allowed to freely explore the three chambers for 5 minutes. In the first testing section, an age-matched stranger mouse (S1) of the same strain was placed in one of the cages, and the subject mouse was allowed to explore the three chambers without restrictions for 10 min. In the second section of preference for social novelty, a second stranger mouse (S2) was placed in the opposite cage, and the subject mouse was allowed to explore again for 10 minutes. All tests were video-recorded, and the time spent in the quadrant around the cage was analyzed using the automated video tracking software TimeCSI (O’Hara & Co., Tokyo, Japan).

### Fear Conditioning Test

Contextual and cued fear conditioning tests were conducted as previously described (43). In brief, on the conditioning trial (Day 1), each mouse was placed in a conditioning chamber (17 × 10 × 10 cm, 100 lux) with clear acrylic walls and a shock-grid floor made of stainless-steel rods (2 mm in diameter, spaced 5 mm apart) and allowed to explore freely for 1 min. A 65-dB tone was then presented for 30 seconds. During the final 2 seconds of the tone presentation, a 0.5-mA foot shock was delivered. The tone and foot shock set were repeated 3 times automatically using a tone and shock generator controlled by Time FZ software (O’Hara & Co., Ltd.). The next day (Day 2), mice were placed in the same chamber without tone or shock for 5 min to evaluate contextual fear memory. On the third day (Day 3), to evaluate cued fear memory, mice were placed in a novel chamber with a dim light (20 lux) and solid white plastic walls without a shock-grid floor and allowed to explore freely for 3 min. The conditioning tone was then presented for 3 minutes. Overhead infrared cameras recorded behavior, and the duration of freezing was automatically calculated using Image FZ2 software.

## Acknowledgements

We thank S. Abe, S. Maeda, H. Eitoku, and T. Morisaki for their technical assistance, K. Tanaka for preparing figures, and D. Hartman for editing the manuscript. We appreciate the technical assistance from The Research Support Center, Research Center for Human Disease Modeling, Kyushu University Graduate School of Medical Sciences, which is partially supported by the Mitsuaki Shiraishi Fund for Basic Medical Research. This work was supported in part by the Leading Program in Fukuoka Bio Valley Project to T.F.; the AMED (JP20gm1310008, JP24wm0625306) and JSPS KAKENHI (JP23H00391) to K.N.; AMED (23gm1310008) and JSPS KAKENHI (JP22K15201, JP23H04170) to H.N.

## Figure Legends

**Extended Data Fig. 1**

**a**, Structural formulas of plasmalogen (top, left) and KIT-13 (top, right). The structures of the other plasmalogen derivatives tested are also shown.

**b**, The body weight of *Mecp2*-KO and KIT-13-treated *Mecp2*-KO mice. Data are represented as mean ± SEM.

**c**, Representative immunofluorescence images of wild-type (WT), *Mecp2*-KO, and KIT-13-treated *Mecp2*-KO mice microglia in the prefrontal cortex. Scale bar = 10 μm.

**d-f**, Quantitative assessment of Iba1-positive microglial cell number (d), soma size (e), and total dendrite length (f). Microglia were measured from n = 3 mice per group. Statistical testing was performed using Tukey’s multiple comparisons test. ^***^*P* <0.001.

