## Supplementary figures and images for "Novel plasmalogen derivative KIT-13 restores neurological function in a mouse model of Rett syndrome by reducing neuroinflammation and restoring mitochondrial function"

### Extended data Fig. 1

**Extended Data Fig. 1**

a

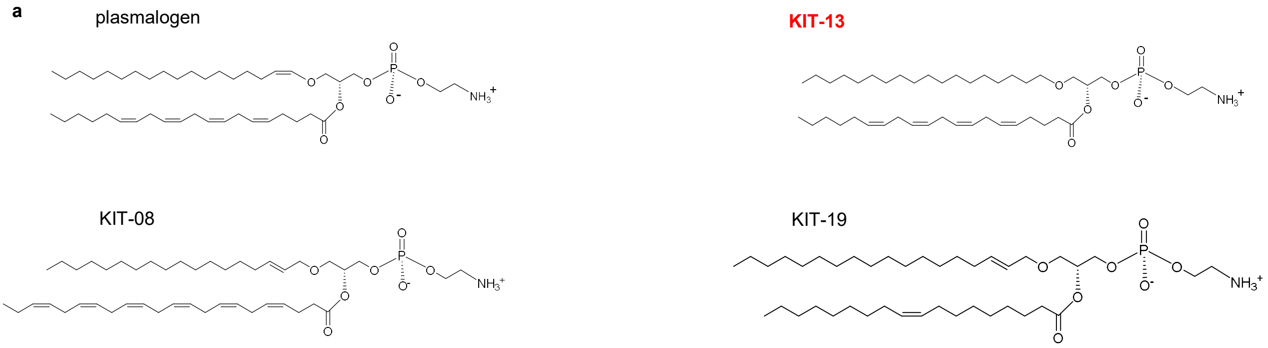


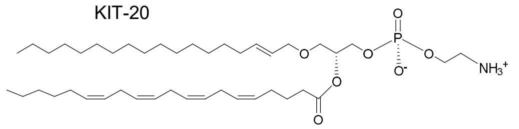


b

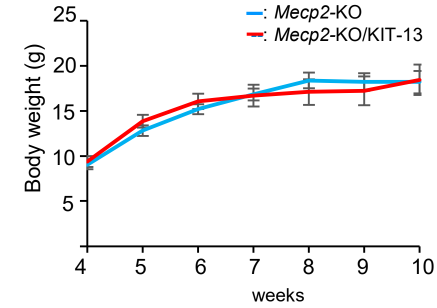


c

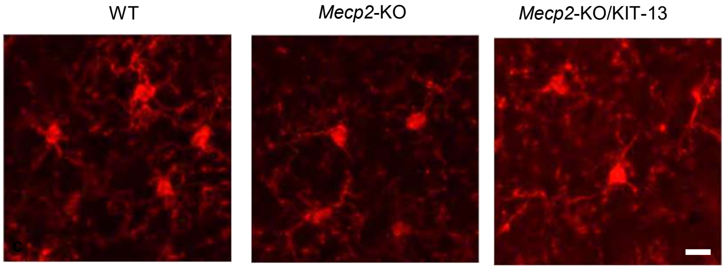


d e f

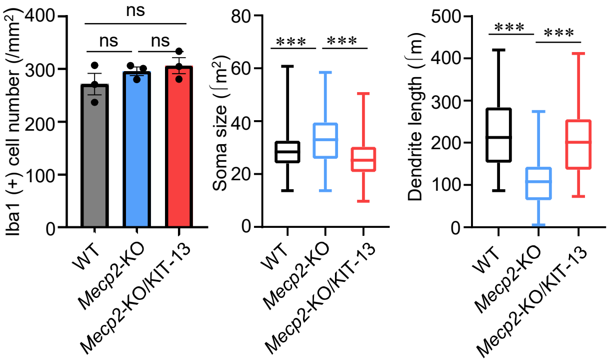
